# Temporal dynamics of inhalation-linked activity across defined subpopulations of mouse olfactory bulb neurons imaged in vivo

**DOI:** 10.1101/558999

**Authors:** Shaina M. Short, Matt Wachowiak

## Abstract

In mammalian olfaction, inhalation drives the temporal patterning of neural activity that underlies early olfactory processing. However, it remains poorly understood how the neural circuits that process incoming olfactory information are engaged in the context of inhalation-linked dynamics. Here, we used artificial inhalation and two-photon calcium imaging to compare the dynamics of activity evoked by odorant inhalation across major cell types of the mouse olfactory bulb (OB). We expressed GCaMP6f or jRGECO1a in mitral and tufted cell subpopulations, olfactory sensory neurons and two major juxtaglomerular interneuron classes, and imaged responses to a single inhalation of odorant. Activity in all cell types was strongly linked to inhalation, and all cell types showed some variance in the latency, rise-times and durations of their inhalation-linked response. Juxtaglomerular interneuron dynamics closely matched that of sensory inputs, while mitral and tufted cells showed the highest diversity in responses, with a range of latencies and durations that could not be accounted for by heterogeneity in sensory input dynamics. Diversity was apparent even among ‘sister’ tufted cells innervating the same glomerulus. Surprisingly, inhalation-linked responses of mitral and tufted cells were highly overlapping and could not be distinguished on the basis of their inhalation-linked dynamics, with the exception of a subpopulation of superficial tufted cells expressing cholecystokinin. Overall, our results support a model in which diversity in inhalation-linked patterning of OB output arises first at the level of sensory input and is enhanced by feedforward inhibition from juxtaglomerular interneurons which differentially impact different subpopulations of OB output neurons.

**SIGNIFICANCE STATEMENT:** Inhalation drives the temporal patterning of neural activity that underlies olfactory processing and rapid odor perception, yet the dynamics of the neural circuit elements mediating this processing are poorly understood. By comparing inhalation-linked dynamics of major olfactory bulb subpopulations, we find that diversity in the timing of neural activation arises at the level of sensory input, which is then mirrored by inhibitory interneurons in the glomerular layer. Temporal diversity is higher among olfactory bulb output neurons, with different subpopulations showing distinct but nonetheless highly overlapping ranges of inhalation-linked dynamics. These results implicate feedforward inhibition by glomerular-layer interneurons in diversifying temporal responses among output neurons, which may be important for generating and shaping timing-based odor representations during natural odor sampling.

## INTRODUCTION

Temporal patterning of neural activity is a fundamental aspect of information coding and processing by neural circuits. In the mammalian olfactory system, the primary driver of temporally patterned activity is inhalation of air through the nasal cavity. Inhalation delivers transient pulses of odorant to the olfactory epithelium, and so determines the initial temporal structure of olfactory sensory input to the brain and drives the temporal patterning of activity at subsequent processing stages (Macrides and Chorover, 1972, Onoda and Mori, 1980, Sobel and Tank, 1993, Kepecs et al., 2006, Schaefer et al., 2006, Schaefer and Margrie, 2007, Wachowiak, 2011). Behavioral and psychophysical studies have shown that odor percepts are formed within the time of a single inhalation (150-250 ms for rodents, ∼400 ms for humans) (Laing, 1986, Johnson et al., 2003, Kepecs et al., 2007), and neurophysiological studies have demonstrated that the temporal pattern of neural activity elicited by a single inhalation of odorant can robustly encode odorant identity and intensity (Uchida and Mainen, 2003, Kepecs et al., 2007, Wesson et al., 2008, Wesson et al., 2009, Cury and Uchida, 2010, Shusterman et al., 2011, Rebello et al., 2014). Thus, understanding how inhalation-linked temporal patterns of activity are generated and shaped by neural circuits in the early olfactory pathway is fundamental to understanding olfactory information processing.

Neural circuits in the olfactory bulb (OB) mediate the first steps in processing olfactory inputs: here, olfactory sensory neurons (OSNs) drive excitation of mitral and tufted cells (MTCs), the principal output neurons of the OB, as well as activate multiple inhibitory circuits within and between OB glomeruli (Wachowiak and Shipley, 2006). This juxtaglomerular inhibition is hypothesized to play a critical role in shaping MTC responses to odorants (Gire and Schoppa, 2009, Shao et al., 2009, Shao et al., 2012, Fukunaga et al., 2014, Banerjee et al., 2015, Liu et al., 2016b) yet the temporal dynamics of activity among different juxtaglomerular interneurons with respect to inhalation have not been well-characterized. Two types of juxtaglomerular interneuron classes – periglomerular (PG) and short axon (SA) cells – are hypothesized to mediate feedforward and lateral inhibition, respectively, with differential impacts on the temporal dynamics of MTC responses (Aungst et al., 2003, Shao et al., 2009, Shirley et al., 2010, Fukunaga et al., 2012, Shao et al., 2012, Liu et al., 2013, Whitesell et al., 2013, Banerjee et al., 2015, Najac et al., 2015, Liu et al., 2016b, Geramita and Urban, 2017). Additionally, feedback inhibition mediated by reciprocal connections between MTCs to PG or SA cells may also shape MTC temporal patterning (Najac et al., 2015).

Here, we sought to better characterize the temporal dynamics of inhalation-driven activity among major circuit elements in the OB in order to refine models of OB circuit function during naturalistic odorant sampling in vivo. We used cell-type specific imaging with GCaMP-based reporters to record from different subpopulations of MTCs as well as presumptive PG cells, SA cells, and OSN inputs. We used an artificial inhalation paradigm to examine responses to a single inhalation of odorant with high fidelity and to precisely compare inhalation-linked response patterns across experiments and between cell types. We found distinct differences in the inhalation-linked temporal patterns of activity among different cell types, with PG and SA cell populations showing faster responses to inhalation than either mitral or tufted cells, and a range of excitatory MTC response dynamics that could not be accounted for by diversity in the dynamics of OSN inputs. At the same time, we observed a great deal of overlap in the inhalation-linked temporal patterns of activity among superficial tufted and mitral cells. Overall, these results support circuit models in which juxtaglomerular interneurons mediate rapid feedforward inhibition that contributes to diverse inhalation-linked temporal patterning in both mitral and tufted cells.

## MATERIALS AND METHODS

### Animals

Genetically-engineered mice expressing Cre recombinase (Cre) targeted to specific neuronal populations were used for experiments. Mice were either crossed to the Ai95 GCaMP6f reporter line (The Jackson Laboratory (JAX), stock #024105) or injected with a viral vector. The mouse strains using included GAD2-IRES-Cre mice (JAX Stock #010802), TH-Cre (JAX stock #008601), DAT-IRES-Cre mice (JAX stock #006660), OMP-Cre (JAX stock #006668), PCdh21-Cre (Gensat stock# 030952-UCD) (Nagai et al., 2005), Tbet-Cre (JAX stock #024507), Thy1-GCaMP6f transgenic mice (JAX stock #024339, line GP5.11) mice, and CCK-IRES-CRE (JAX stock #012706). Mice were on average 4.6 months of age by completion of data collection. Both female (46) and male (54) mice were used. Mice were housed up to 5 per cage in a 12/12h light/dark cycle. Food and water were provided *ad libitum.* Each procedure was performed following the National Institutes of Health Guide for the Care and Use of Laboratory Animals and approved by the University of Utah institutional animal care and use committee.

### Viral vector expression

Cre-dependent expression of GCaMP6f (AAV2/9, AAV2/1, or AAV2/5 stereotypes of hSyn.Flex.GCaMP6f) or jRGECO1a (AAV2/9 or AAV2/5 hSyn.NES.jRGECO1a.WPRE.SV40) was achieved with the injection of viral vectors. All viruses were obtained from the University of Pennsylvania Viral Vector Core. Injections were made into the dorsal OB under isoflurane anesthesia (0.5-2% in O_2_). Where applicable, mice used for virus injection were homozygous for the allele driving Cre expression. Using a stereotaxic head holder and drill, a small craniotomy (0.5-1mm) was made on the dorsal surface of the OB. A glass pipette was lowered to a depth 50-150 µm to target periglomerular or short axon interneurons or 200-400 µm to target mitral and tufted cell populations. Mice were single housed following the surgery and imaged 14-28 d after injection.

### In vivo imaging

Two-photon imaging was performed in acutely anesthetized mice. Initially, pentobarbital (50 mg/kg) was used during the implantation of a double tracheotomy (Wachowiak and Cohen, 2001, Bozza et al., 2004, Spors et al., 2006), after which isoflurane (0.5 −2% in O_2_) was delivered directly to the tracheotomy tube, bypassing the nose. Next, a custom head bar was implanted, a craniotomy was made, and a coverslip was implanted using 2.5% low-melting-point agarose over the dorsal OB for imaging (Wachowiak et al., 2013). Throughout surgeries and while imaging, the body temperature was maintained at 37°C with a heating pad and the heart rate at ∼400 beats per minute.

Imaging data were collected using either a Moveable Optical Microscope (Sutter Instruments) coupled to a Mai Tai HP pulsed Ti: Sapphire laser (Newport Corp.) and controlled by Scanimage 5.1 (Pologruto et al., 2003), or a Neurolabware microscope coupled to a Cameleon UltraII laser (Coherent) and controlled by Scanbox software. Both setups used resonance-based scanning and GaASP photomultipliers (Hamamatsu H10770B) for light collection, and images were collected at a frame rate of 15.5 Hz. A 16X 0.8 N.A. (Nikon) objective was used in all experiments. For dual-color imaging, a Fidelity 1070 nm femtosecond laser was used simultaneous with 920 nm illumination and emission filters were used to separate green (520/65 nm) and red (641/75 nm) emission (Sun et al., 2017).

### Analysis of imaging data

Maps of inhalation-triggered fluorescence changes (i.e., ITA response maps) were generated by choosing 15 frames before and after odorant inhalation. For display, ITA response maps were smoothed with a Gaussian filter with sigma 1.25 pixels. Regions of interest (ROIs) were selected manually from ITA response maps or from resting fluorescence images. Fluorescence time series were extracted by averaging all pixels in a ROI using custom MATLAB scripts. All time series data was sampled to 150 Hz using the Matlab piecewise cubic interpolation functions interp1 and pchip. In all cases, ΔF/F was calculated as (ΔF/F=(F-F_o_)/F_o_), with F_o_ being the mean fluorescence prior to the inhalation, averaged for each inhalation. However, for population level analyses, signals were averaged across three trials of 20s odorant presentations (Fig 1). Excitatory events were defined as ITA responses that were greater than 4 standard deviations (SD) above the ITA baseline signal, which was defined as 1 second prior to inhalation, whereas inhibitory events were defined as ITA responses reaching more than 3 SD below baseline.

**Figure 1.**
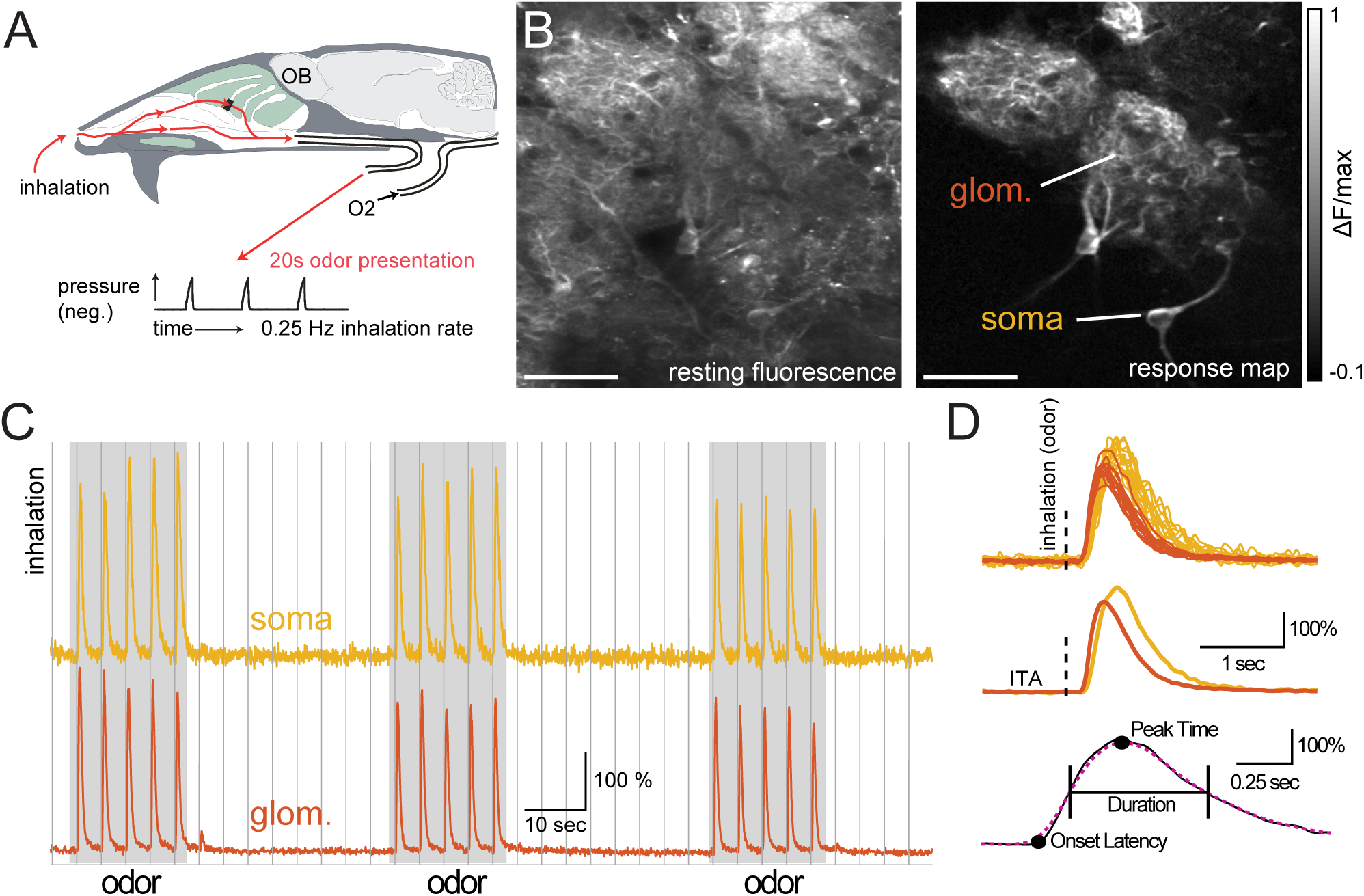
Imaging inhalation triggered temporal dynamics of OB neuron subtypes. (A.) Schematic of preparation for artificial inhalation in the anesthetized mouse. Inhalations were generated by a vacuum pulse applied to nasopharynx at 0.25 Hz. (B.) Example resting fluorescence (left) and inhalation-triggered average (ITA) evoked ΔF/F map of responses taken from a glomerulus (glom.) and interneuron somata (soma) in the glomerular layer in a GAD2-Cre:GCaMP6f reporter cross. White scale bar = 50 µm. Presumptive PG cell somata and glomerular boundaries are difficult to distinguish in the resting fluorescence image, but are clearly apparent in the response map. (C.) Traces showing the fluorescence signal imaged from the glomerular neuropil and soma of an associated GAD2+ neuron, from the example in (B). Traces show a typical imaging epoch, with three 20s odor stimuli (grey) presented during 0.25 Hz inhalation (vertical lines). (D.) Overlaid inhalation triggered calcium signals (top) and corresponding ITAs (middle) taken from the glomerulus and soma during each inhalation in the presence of odorant. Dashed line indicates inhalation onset. Bottom: Schematic illustrating definition of onset latency, time to peak, and response duration. Onset latency was calculated from the ITA trace (solid black line) when the response initially surpassed at least 4 consecutive frames, 4 standard deviations (SD) above baseline. Time to peak (time to maximum response) and response duration (time between the half max and half min points) were calculated from the ITA after applying the Gaussian weighted box filter (dotted magenta line).

### Analysis of temporal dynamics

Onset latency was calculated as the first time point in which the following 4 frames of an ITA trace were above the threshold for a significant excitatory response (4 SD above baseline, which was taken from a 1 s pre-stimulus window). Peak response amplitudes and time to peak values were calculated from ITAs that were filtered using a Gaussian-weighted moving average filter with a window length of 270 ms. From this filtered trace, response duration was calculated as the time from 50% of peak response on the rising slope of the signal to 50% of peak on the decaying slope of the signal. Time to peak was calculated from the maximum value of the filtered trace. For display of pseudo-color plots of ITA responses (e.g., Fig. 2C), the mean of the 1s pre-stimulus window was subtracted from the response time series data, each signal was normalized to its own max and negative max amplitudes, and a Gaussian boxcar filter was applied using a moving average window of 100ms before plotting. For comparisons of glomerular neuropil dynamics to PG and sTC somatic responses (e.g., Fig. 7), values measured from the glomerular ITA were subtracted from those of the somatic ITAs and the variance of these responses calculated using the standard Matlab variance function, which takes the sum of the squared difference between each these somatic signal and the mean of all somatic signals, times 1 over the number of observations minus 1.

**Figure 2.**
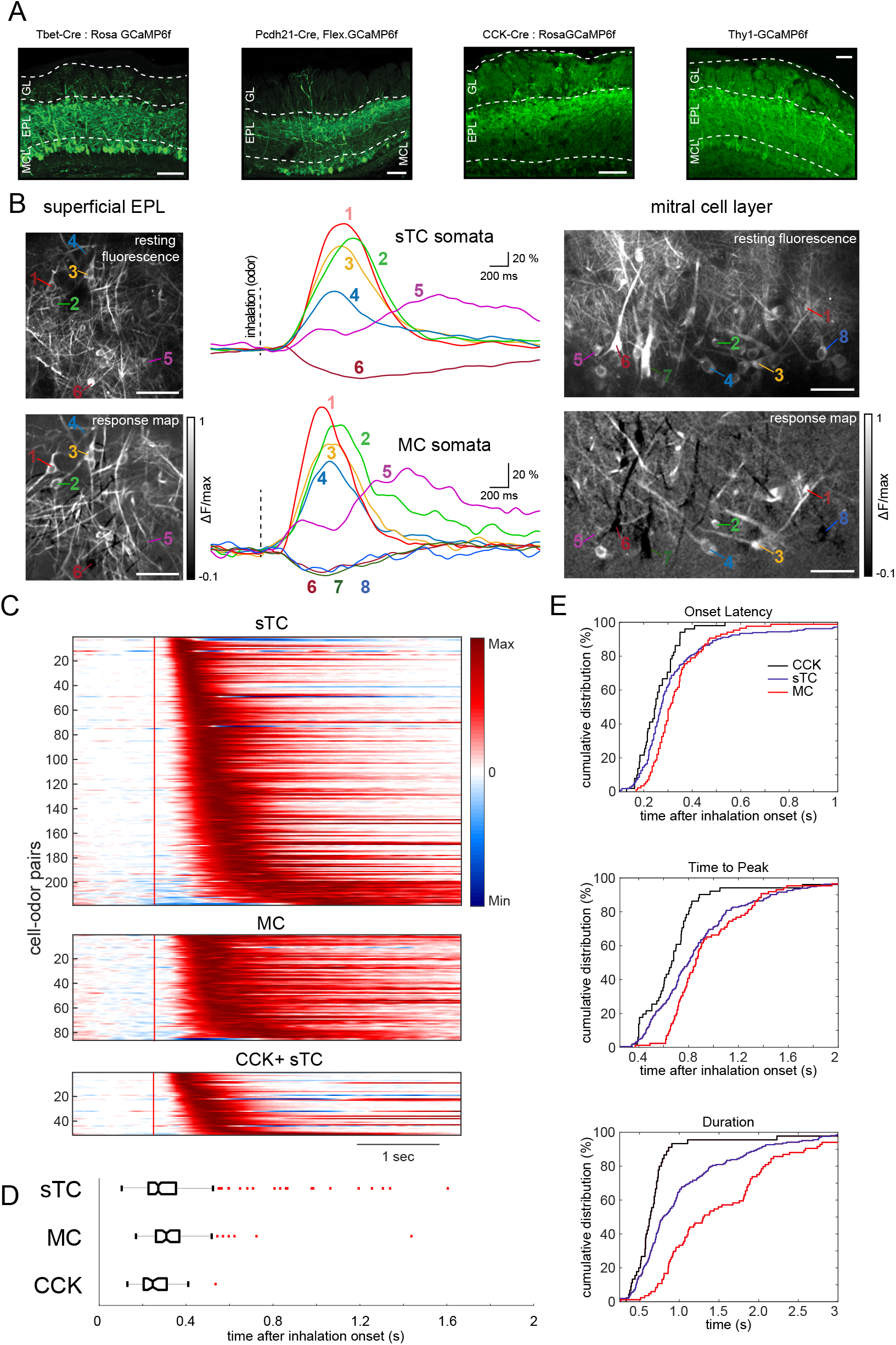
Inhalation-linked dynamics of mitral and superficial tufted cells are diverse and highly overlapping. A. GCaMP6f expression targeted to MC and sTC populations using reporter crosses to target GCaMP6f expression in Tbx21+, CCK+, and Thy1+ MTC populations and virus injections into the olfactory bulb to target GCaMP6f expression to PCDdh21+ MTCs. Glomerular layer (GL), external plexiform layer (EPL), and mitral cell layer (MCL) are labeled using dotted white lines. B. Resting fluorescence (top) and ITA response maps (evoked ΔF/F map, bottom) of sTCs imaged from superficial EPL (left images) and MCs imaged from the mitral cell layer (right images). Middle column shows ITA traces from sTCs (top) and MCs (bottom) indicated in the images. Signals were low-pass filtered at 3 Hz prior to averaging and scaled relative to the pre-inhalation baseline (i.e., ΔF/F). EPL and MCL images were taken from fields of view that were immediately above and below each other in the same Tbet-Cre: Rosa-GCaMP6f mouse during stimulation with the same odorant (methyl valerate). All white scale bars = 50 µm. C. Pseudo-color plots of ITA responses for all excitatory responses in sTC (top), MC (middle), and CCK+ sTCs (bottom); each row shows a different cell-odor pair. Each trace (row) was normalized to its own max and negative max amplitudes on a scale from −1 to 1. Red vertical line indicates inhalation onset. D. Box and whisker plots (top panel) of sTC, MC, and CCK+ sTC onset latency distributions. Outliers marked with red horizontal dashes.(E) Cumulative distribution plots of onset latency (top), time to peak (middle), and duration (bottom) for MC, sTCs, and CCK+ sTC ITAs.

### Experimental Design and Statistical Tests

All statistical details of experiments are listed in the Results section. All datasets (onset latency, time to peak, and response durations across all sTC, MC, OSN, PG, and SA cell populations) rejected the null hypothesis for one-sample Kolmogorov-Smirnov test for normality, therefore non parametric statistical tests were performed as stated throughout the methods. A Wilcoxon signed rank test tested was used for paired comparisons across two groups (mitral and tufted cell populations Fig. 3), Mann-Whitney U-test was used for unpaired comparisons across two groups (mitral and tufted cell population Fig. 2), and the Kruskal-Wallis test was used for comparisons across all subpopulation (Fig. 6). Post hoc multiple comparisons were performed using Tukey’s honest significant difference criterion (Fig. 6). Statistical significance was set at p<0.05. All of these statistical tests were performed in Matlab.

**Figure 3.**
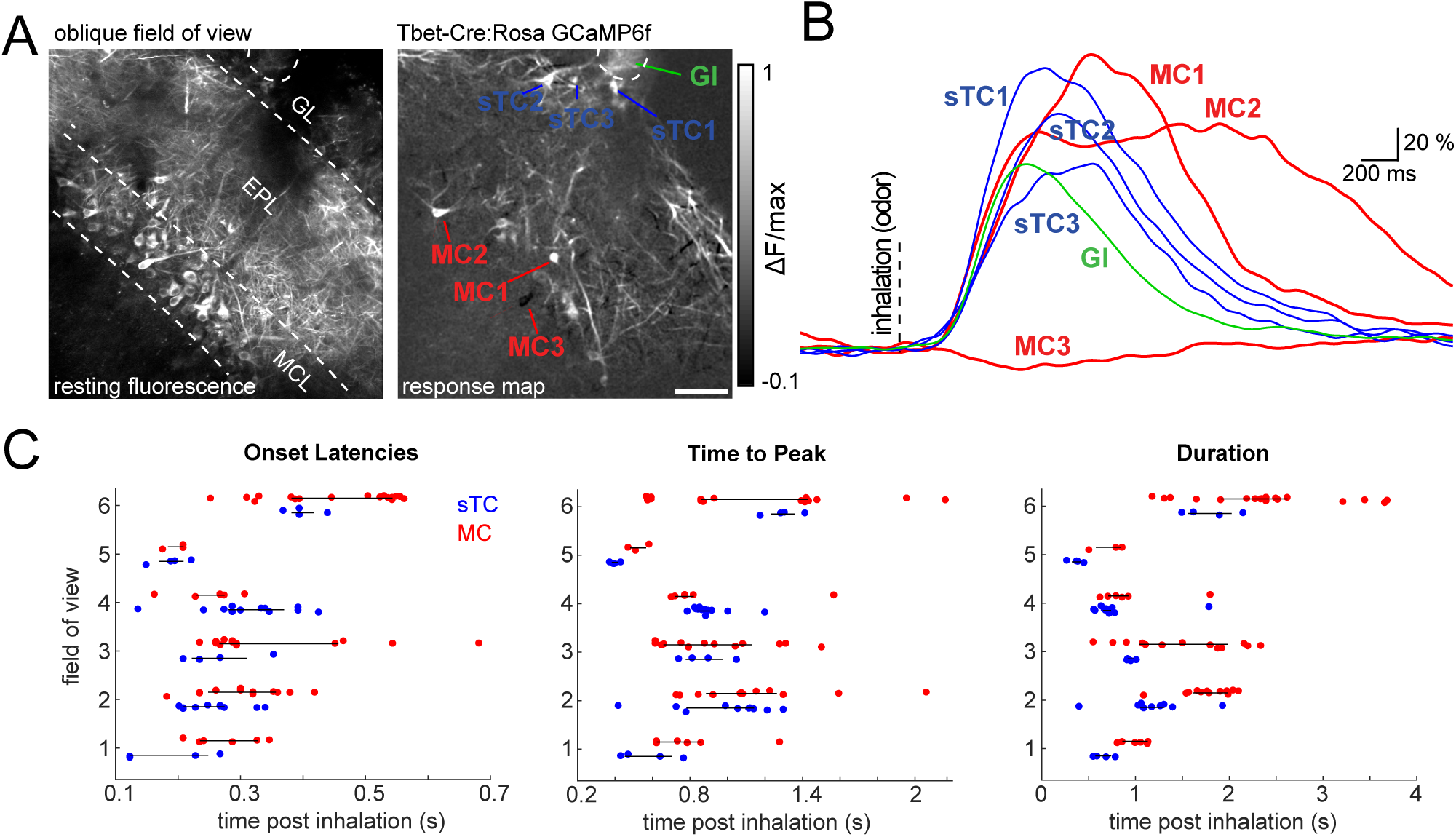
Diversity of inhalation-linked dynamics of mitral and superficial tufted cells within the same field of view. A. Resting fluorescence and ITA response map showing sTCs and MCs imaged in the same field of view using an oblique imaging plane in a Tbet-Cre: Rosa-GCaMP6f mouse during stimulation with methyl valerate. Gl, glomerular neuropil. B. Traces showing ITAs from the sTCs and MCs shown in (A), illustrating temporal diversity of inhalation-linked responses across different cell types imaged simultaneously. C. Plots comparing latency, peak time and duration of ITAs from sTCs (blue) and MCs (red) imaged in the same field of view or successive z-planes, allowing for paired comparisons (see Text). Black horizontal line spans the interquartile range. White scale bars, 50 µm.

### Olfactometry

A custom olfactometer controlled by Labview software was used to present odorants, as previously described (Bozza et al., 2004, Verhagen et al., 2007). Odorant concentration was controlled by diluting from saturated vapor in filtered, ultra-pure air. Odorants were present for 20 sec at 1-5% saturated vapor. Each odorant presentation was separated by a 32s inter-stimulus interval. Odorants were obtained at 95-99% purity (Sigma-Aldrich) and stored under nitrogen. Some odorants were diluted in mineral oil to achieve final concentrations at the animal’s nose of 1-20 ppm at 1% saturated vapor. Odorants tested included ethyl butyrate, methyl valerate, butyl acetate, 2-hexanone, ethyl tiglate, 2-methyl pentanal, 2-hydroxyacetophenone, and hexyl acetate. Odorants were delivered 0.5 – 1.0 cm in front of the mouse’s nose. Filtered, ultra-pure air was delivered to the mouse’s nose in between odorant presentations. A fan positioned behind the animal scavenged excess odorant in the room.

### Histology

An overdose of sodium pentobarbital was used to deeply anesthetize mice prior to perfusion with phosphate-buffered saline (PBS) followed by 4% paraformaldehyde in PBS. Overnight, heads were post fixed in 4% paraformaldehyde in PBS. Next, the brain was extracted and embedded in 5% agarose. Coronal sections (100-200 µm thick) were made with a Vibratome and mounted onto a glass coverslip before imaging with a Fluoview FV1000 Olympus confocal microscope at 10X, 20X, and 40X magnification.

## RESULTS

To compare the dynamics of inhalation-driven activity across different OB neuron populations, we used an artificial inhalation paradigm that allowed for precise comparison of response dynamics across experiments, as described previously (Wachowiak et al., 2013, Diaz-Quesada et al., 2018) (Fig. 1 A). We generated single inhalations at 0.25 Hz to enable inhalation-triggered averaging of responses with, in most cases, minimal adaptation from one inhalation to the next. We expressed calcium sensors (typically GCaMP6f) in distinct genetically-defined cell types using either viral vectors or genetic expression strategies, and imaged activity from the somata of the targeted cell types or from glomerular neuropil using two-photon laser scanning microscopy (Fig. 1B). A typical imaging epoch consisted of three periods of odorant presentation, each lasting 20 seconds (i.e., 5 inhalations at 0.25 Hz), with a 32 sec interval between presentations (Fig. 1C), yielding a total of 15 inhalations in the presence of odorant and 10 inhalations of clean air. We analyzed response dynamics from inhalation-triggered average traces (ITAs) and determined response magnitudes, latencies, and durations as we have done previously (Carey et al., 2009)(Fig. 1D).

### Inhalation-linked dynamics of mitral/tufted cell subpopulations

We first examined inhalation-linked response dynamics in mitral and tufted cells (MTCs). Electrophysiological recordings have shown diverse inhalation-linked temporal patterns among MTCs and distinct differences between these subpopulations as defined by soma depth (Fukunaga et al., 2012, Igarashi et al., 2012). To test if these differences were reflected in inhalation-linked calcium signals, we selectively expressed GCaMP6f in MTCs via several mechanisms: viral injection (AAV.Flex.GCaMP6f) into the OB of mice expressing Cre-recombinase in protocadherin-21 positive (PCdh21-Cre) neurons (Wachowiak et al., 2013); crossing a Cre-dependent GCaMP6f reporter line with mice expressing Cre in Tbx21-positive (Tbet-Cre) neurons (Haddad et al., 2013) or in cholecystokinin-positive (CCK+) neurons (Seroogy et al., 1985); and use of a transgenic mouse line (Thy1-GCaMP6f) selective for expression in MTCs (Dana et al., 2014). Patterns of GCaMP6f expression using PCdh21-Cre, Tbet-Cre, or Thy1-GCaMP6f mice were qualitatively similar, as described previously (Wachowiak et al., 2013) with expression in large numbers of MTCs (Fig. 2A). To compare mitral and tufted cells, we distinguished each population by somatic depth (Fig. 2B). We restricted our analysis to somata that were clearly in the mitral cell layer (MCs) and superficial tufted cells (sTCs) just below the glomerular layer, excluding deeper or middle tufted cells in the external plexiform layer.

For both MCs and sTCs, the predominant inhalation-linked response pattern was a transient fluorescence increase, presumably corresponding to a brief spike burst after inhalation. There was substantial diversity in the dynamics of this transient, with different cells showing differences in onset latency, time to peak response, and response duration; such diversity was apparent for different cells imaged within the same field of view (Fig. 3C). While MCs and sTCs showed statistically significant differences in latency, time to peak, and response duration (p = 0.0032, 0.0119 and 3.3e^−9^ respectively, Mann-Whitney U test, MC=86 cell-odor pairs from 5 mice, sTC=214 cell-odor pairs from 16 mice), their response dynamics were highly overlapping (Fig. 2C-D). MCs and sTCs differed most substantially in their response durations, with MCs showing significantly longer-duration responses (median, 1309 ms, interquartile range: 892 – 1994 ms) compared to sTCs (median, 788 ms, interquartile range: 611 – 1296 ms, Fig. 2E). Another difference observed between sTCs and MCs was their responsiveness to inhalation of clean air, with 9.5% of sTCs (22/231 cells) and zero MCs (0/89 cells) showing a significant response.

In several cases we were able to more directly compare sTC and MC response dynamics, either by imaging sTCs and MCs in the same field of view using an oblique imaging plane (n = 3 fields of view from 2 mice) (Fig. 3A) or by imaging sTC and MC responses to the same odorant in successive trials by shifting the focal plane from the superficial external plexiform layer to the mitral cell layer (n = 3 paired imaging planes from 2 mice). Even with this within-preparation comparison, sTC and MC response dynamics were still highly overlapping (Fig. 3B). There was no significant difference in the median ITA onset latency when comparing MC and sTC populations within the same preparation or field of view (Fig. 2C, left panel, Wilcoxon signed rank test, p=0.16, n = 6 paired comparisons, 58 total MCs, 38 total sTCs). Likewise, there was no significant difference in median sTC and MC ITA time to peak (Fig. 3C, middle panel, Wilcoxon signed rank test p=0.22). However, the median half width durations of sTC responses were significantly shorter compared to MC responses (Fig. 3C, right panel, Wilcoxon signed rank test p=0.03). This analysis supports the conclusion that, as a population, sTCs show slightly shorter-duration responses than MCs, but that individual sTCs and MCs overlap substantially in their inhalation-linked response dynamics.

Finally, we measured inhalation-linked responses in sTCs defined by their expression of the peptide transmitter cholecystokinin (CCK), which likely constitute a subset of Tbx21+, Thy1+, or PCdh21+ sTCs (Seroogy et al., 1985, Liu and Shipley, 1994, Tobin et al., 2010) and which we have previously shown to exhibit shorter-onset and simpler odorant-evoked responses than MCs (Economo et al., 2016). Consistent with these earlier reports, sTCs imaged from GCaMP6f:CCK-Cre mice (n= 52 cell-odor pairs in 3 mice) indeed showed onset latencies that were significantly shorter than those of MCs (p = 3.7e-5) or the general sTC population (p = 0.0333) defined by Tbx21, PCdh21, or Thy1 expression (median onset latency: 247 ms, interquartile range: 206-311 ms; Mann-Whitney test, Fig. 2C-E). Likewise, CCK+ sTCs showed an earlier time-to-peak than the general sTC population (p = 0.0022, Mann-Whitney test). CCK+ sTCs showed the largest difference in their response durations, which were uniformly short (median, 635 ms, range: 520 – 735 ms) and significantly shorter than the general sTC population (p = 9.1e^−5^, Mann-Whitney test). Thus, CCK+ sTCs appear to constitute a distinct subpopulation of sTCs with particularly rapid inhalation-triggered response patterns.

We also examined inhalation-linked suppression in MTCs. We and others have previously reported that about a third of MTCs show odorant-evoked suppression of ongoing activity (Kollo et al., 2014, Economo et al., 2016, Diaz-Quesada et al., 2018). We assessed whether phasic suppression elicited by each inhalation was apparent in GCaMP6f signals, using a conservative criterion of a fluorescence decrease in the ITA of at least 3 SD below the pre-inhalation baseline. Using these criteria, odorant-evoked, inhalation-linked suppression was sparsely distributed and relatively rare (Fig. 4A), with only 8 of 93 (9%) MC-odor pairs and 5 of 240 (2%) sTCs showing suppressive ITAs. This prevalence is substantially smaller than the prevalence of suppressive responses seen in awake, freely-breathing mice or during higher-frequency (2 Hz) artificial inhalation (Economo et al., 2016), or as measured with whole-cell recordings (Kollo et al., 2014, Diaz-Quesada et al., 2018). A possible explanation for this difference is that the ability to detect inhalation-linked suppression using the GCaMP6f reporter was clearly dependent on baseline activity levels in individual cells, which could fluctuate over the course of a trial (Fig. 4A). Notably, both MCs and sTCs could also show phasic suppression linked to inhalation of clean air alone (example sTC Fig. 4B), with such suppression more prevalent among sTCs than MCs (21 of 231 sTCs versus 2 of 89 MCs). Even in these cases, however, inhalation-linked suppression was sparsely distributed among the multiple cells in a field of view (Fig. 4B).

**Figure 4.**
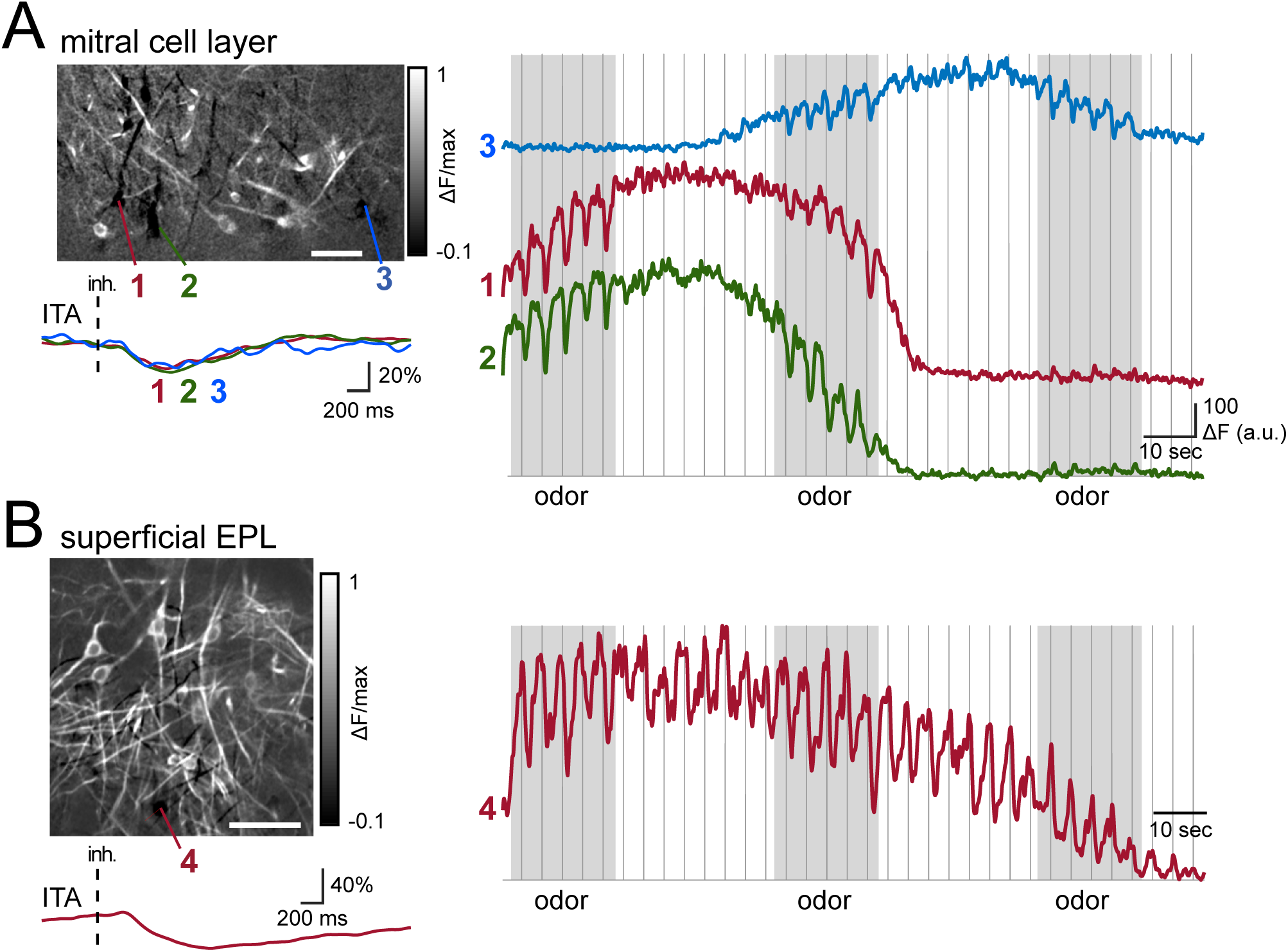
Inhalation-linked suppression in mitral and superficial tufted cells. A. Left: Odorant-evoked ITA response map (evoked ΔF/F) including several MCs (Tbet-Cre: Rosa-GCaMP6f mouse) that show suppression of ongoing activity after each inhalation of odorant (methyl valerate, same preparation shown in Fig. 2B), with ITA traces from each cell shown below. A 3 Hz low pass filter was applied to all traces prior to averaging. Right: Continuous traces from each MC showing fluorescence decrease after odorant inhalation which is only apparent when ongoing activity (reflected in pre-odor fluorescence) is sufficiently high. B. Same as in (A) but showing responses of sTCs imaged from the superficial EPL in the same animal in (A). The field of view is directly above that shown in (A). The odorant is also the same as that in (A). Trace shows ITA and continuous recording from a suppressed sTC. This cell was suppressed by inhalation of clean air with no change during odorant presentation. All white scale bars = 50 µm.

Overall, these data suggest that the chief difference in the inhalation-triggered dynamics of MCs versus sTCs is that MCs show a greater range of excitatory response latencies and durations than do sTCs. At the same time, they also suggest that sTC and MC responses do not unambiguously map to any single parameter of the inhalation-linked response, including response latency, response duration, or even response polarity. We next used this same approach to gain insight into where in the OB circuit the diversity in response patterns might arise by imaging inhalation-triggered responses from olfactory sensory neurons (OSNs) and juxtaglomerular interneurons.

### Contribution of olfactory sensory input dynamics to MTC response diversity

One determinant of diverse MT cell inhalation-linked response dynamics could be diversity in the temporal patterns of sensory input to the OB (Spors et al., 2006). To assess this we measured the temporal dynamics of OSN input to OB glomeruli using GCaMP6f expressed in OSNs (Fig 5A) and imaging responses from OSN axon terminals, as described previously (Wachowiak et al., 2013). Consistent with earlier studies (Spors et al., 2006, Carey et al., 2009, Wachowiak et al., 2013), OSN responses were predominately simple transient fluorescence increases following each inhalation (Fig 5B-D). Surprisingly, inhalation of clean air elicited significant excitatory responses in only 1 of 72 glomeruli imaged (1/72), a lower fraction than expected given prior reports of inhalation-linked excitation among OSNs (Grosmaitre et al., 2007, Carey et al., 2009). We also observed inhalation-linked suppressive responses in a small fraction (3/72) of glomeruli imaged.

**Figure 5.**
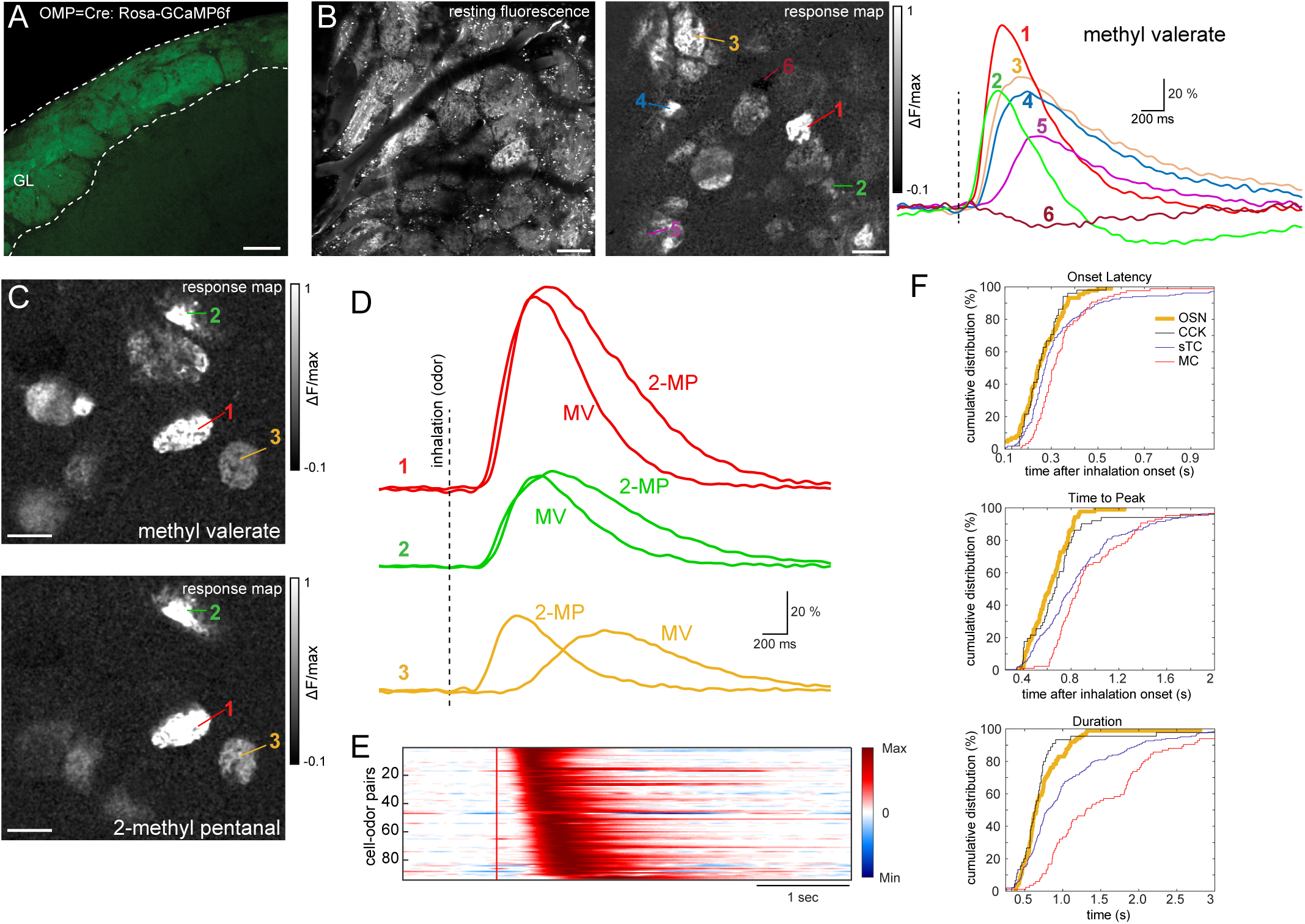
Comparison of inhalation-linked dynamics of OSN inputs and MTCs. A. GCaMP6f expression in OSN axons innervating OB glomeruli as seen in confocal histology in an OMP-Cre:Rosa-GCaMP6f mouse. B. In vivo resting fluorescence (left) and odorant-evoked ITA response map imaged across glomeruli of the dorsal OB. ITA response traces from the indicated glomeruli are shown at right. Note the range of excitatory dynamics and the presence of a suppressive response in one glomerulus. C. ITA response maps (left) and traces (right) showing three glomeruli that each respond to two odorants: methyl valerate (MV) and 2-methyl pentanal (2-MP). ITA traces for each odorant are overlaid for the same glomerulus, illustrating odorant-specific response dynamics. D. Pseudocolor plots of ITA responses for all odor-glomerulus pairs, displayed and normalized, as in Fig. 2, to their minimum (−1) and maximum (1) amplitude response. Red vertical line indicates inhalation onset. E. Cumulative distribution plots showing OSN ITA onset latencies, times to peak, and durations, with the values for MCs, sTCs, and CCK+ sTCs (same data as in Fig 2) included for comparison. All scale bars, 100 µm.

With respect to odorant-evoked activity, OSN ITAs varied in their latency of onset, rise-time and response duration, and different odorants could elicit different inhalation-triggered dynamics within the same glomerulus (Fig. 4C, D). As a population, OSN response onset latencies were significantly earlier than those of MCs, with 50% of glomerular input latencies preceding the shortest 82.6% of MC responses (Fig. 4E, F, median, 242 ms, range, 197-321 ms, n= 88 glomerulus-odor pairs, 5 mice). Similar differences in temporal dynamics between OSN and MTC responses appeared for time to peak and response duration, with OSN responses being earlier and shorter than those of MCs. The largest difference between OSN and MTC ITA dynamics was in the duration of the OSN versus MC responses, with half of all MCs showing ITA durations longer than 72.7% of all OSN responses. Notably, the distributions of onset latencies, rise-times, and durations for OSN inputs overlapped closely with those of CCK+ sTCs (Fig. 4F). These results are consistent with a model in which inhalation-linked excitatory responses among sTCs – and in particular, CCK+ sTCs-largely reflect excitatory drive from OSNs, while MC excitation is further shaped by additional synaptic or intrinsic mechanisms (Kikuta et al., 2013, Adam et al., 2014).

### Temporal dynamics of inhalation-linked activity in juxtaglomerular interneurons

We next characterized inhalation-linked responses in juxtaglomerular interneurons, focusing on periglomerular (PG) and short axon (SA) cells – these two classes of are thought to shape MTC responses via feedforward and lateral inhibition (Kosaka and Kosaka, 2008, Liu et al., 2013, Fukunaga et al., 2014, Banerjee et al., 2015, Liu et al., 2016a). GCaMP6f was preferentially targeted to PG or SA cells by either virus injection (AAV.Flex.GCaMP6f) or Rosa-GCaMP6f reporter cross using GAD2-Cre mice (for PG cells) or TH-Cre or DAT-Cre mice (for SA cells), as described previously (Wachowiak et al., 2013, Banerjee et al., 2015) (Fig. 6A, D). Calcium signals were imaged from somata located around the glomerulus periphery (Fig. 6B, E).

**Figure 6.**
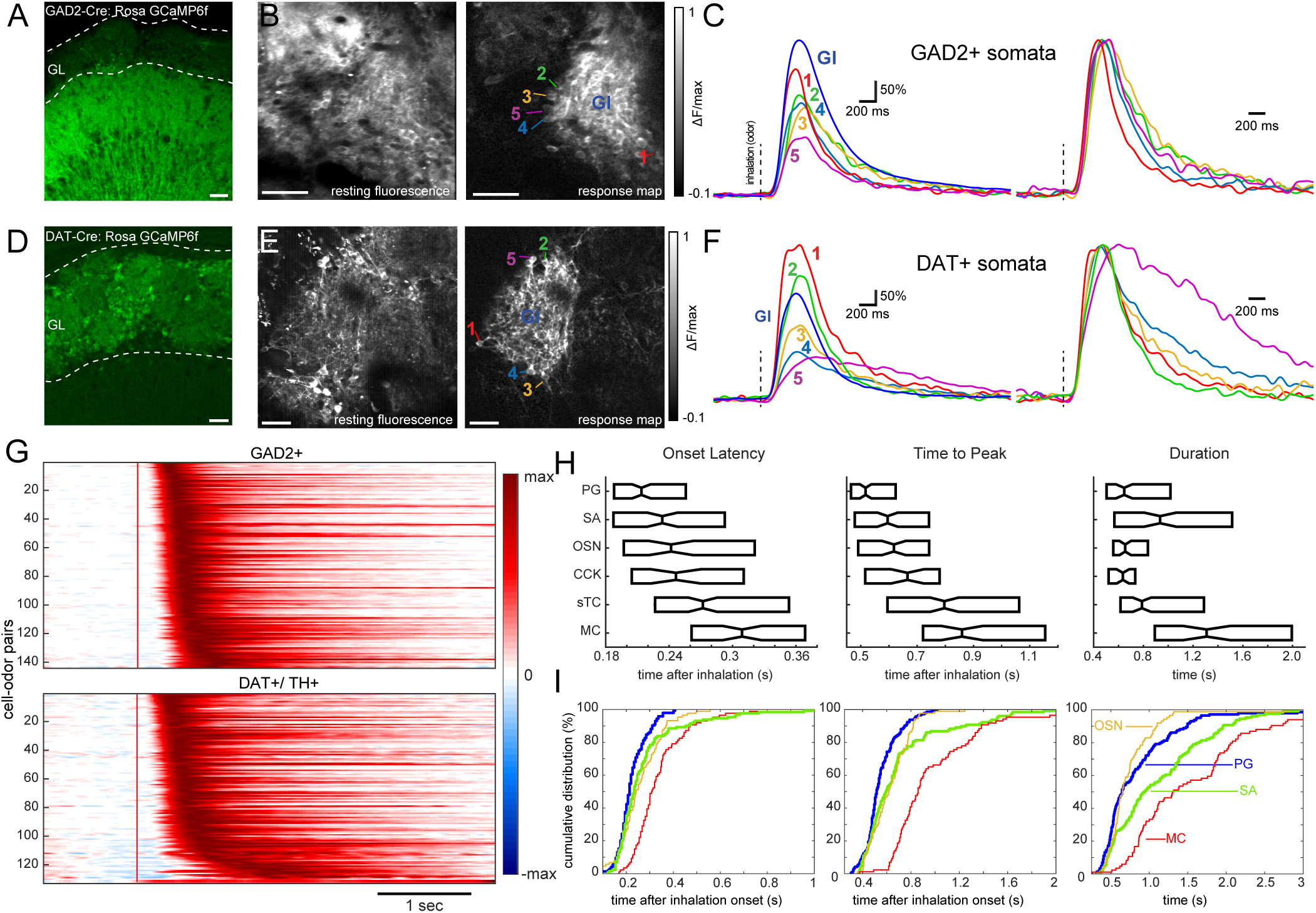
Juxtaglomerular interneurons show simple and short-latency responses to odorant inhalation. A. GCaMP6f expression in the glomerular layer of a GAD2-IRES-Cre mouse, showing expression in juxtaglomerular neurons with extensive processes in the glomerular layer (GL). Scale bar, 50 µm. B. In vivo resting fluorescence (left) and odorant-evoked ITA response map (right) showing activation of two adjacent glomeruli in a GAD2-Cre: Rosa GCaMP6f mouse. Note that numerous somata are apparent around the periphery of the right glomerulus in the response map. Numbers indicate somata whose ITAs are shown in (C). Gl, glomerular neuropil. Scale bar, 50 µm. Odorant was methyl valerate. C. ITA traces from the somata and glomerular neuropil from (B). Traces were low-pass filtered at 5Hz prior to averaging. The same traces are shown scaled to the same peak values at right. Note that all cells show similar onset latencies with only modest differences in time to peak or duration. D-F. Similar to (A-C), but for dopaminergic neurons using a DAT-Cre: Rosa GCaMP6f mouse to target presumptive SA cells. The ITA traces for these DAT+ cells are also all short-latency, with one cell showing a longer-lasting response. Odorant was 2-Hexanone. G. Pseudocolor plots of ITA responses for all cell-odor pairs for GAD2+/presumptive PG cells (top) and DAT+/TH+ presumptive SA cells (bottom), displayed and normalized as in Fig. 2. Red vertical line indicates inhalation onset. H. Box plot comparisons of ITA onset latencies, time to peak, and half width durations (medians (center horizontal line) ± 2^nd^ and 3^rd^ quartiles across all cell populations investigated. (I) Cumulative distribution plots of onset latencies, times to peak, and durations for GAD2+ and DAT/TH+ cells highlighted by thicker plots. OSN and MTC populations shown for reference.

Both GAD2+ PG and TH/DAT+ SA cell ITA odor responses consisted overwhelmingly of simple, monophasic response transients; multiphasic responses were not seen (Fig. 6C, F, G). In contrast to MCs and sTCs, odorants did not elicit suppressive responses in PG (1/145 cell-odor pairs) or SA cells (0/132 cell-odor pairs), although clean air did elicit excitatory responses in a fraction of both cell types (7/136 PG cells; 2/120 SA cells). With respect to odorant-evoked response dynamics, PG and SA cell populations both had short-latency ITAs, with the main difference being a ‘tail’ of longer-latency SA cell responses (Fig. 6G, H, J). Indeed, the median PG cell ITA latency preceded that of 94% of all MCs, whereas the median SA cell ITA latency preceded 88% of all MCs. At the population level, OSN inputs, PG, and SA cells all had ITA onset latencies that were statistically similar to each other and distinct from that of MCs and other sTCs (Fig. 6IJ, left panels, Kruskal-Wallis test, Chi-sq=100.45, df=715, P>Chi-sq=4.2e^−20^, tukey-kramer post hoc test, p<0.05). Similar trends were observed when comparing time to peak response across these populations (Fig 6IJ, middle panels, Kruskal-Wallis test, Chi-sq=162.05, df=715, P>Chi-sq=3.6e^−33^, tukey-kramer post hoc test, p<0.05).

SA cell responses differed from those of PG cells mainly in their response durations (Fig. 6IJ, right panels, PG half width median, 646 ms, interquartile range: 504 – 1018 ms, SA median, 935 ms, interquartile range: 566 – 1514 ms), with SA cell durations shifted towards longer values than those of PG cells (Fig. 6IJ, right panels). Indeed, there was no significant difference in response duration of OSNs, PG cells, and CCK+ sTCs, while SA cell response durations were longer and statistically indistinguishable from that of other sTCs (Kruskal-Wallis test, Chi-sq=104.62, df=698, P>Chi-sq=5.6e^−21^, tukey-kramer post hoc p<0.05). Overall, these results suggest that inhalation-linked PG and SA responses largely follow those of OSN inputs, while a subset of SA cells exhibit longer-lasting responses. Their short onset latencies are consistent with both cell types mediating rapid feedforward inhibition of MCs and sTCs during inhalation.

### The diversity of inhalation-linked response dynamics within a single glomerulus

PG cells, sTCs, and MCs all receive excitatory input from dendrites confined to a single glomerulus, and there is evidence that different MCs associated with the same glomerulus (i.e., sister MCs) can show distinct temporal response patterns (Dhawale et al., 2010, Arneodo et al., 2018), while sister PG cells have been reported to show temporally uniform response latencies (Homma et al., 2019). We thus next asked to what degree does variance in inhalation-linked response patterns reflect heterogeneity among sister PG cells or sTCs, as opposed to simply reflecting glomerulus-and odorant-specific diversity in OSN input dynamics (Spors et al., 2006).

Initially we directly compared PG and sTC response dynamics during the same odor stimulus and within the same glomerulus using two-color imaging. We used Thy1-GCaMP6f: GAD2-Cre crosses, expressing the red shifted calcium indicator jRGECO1a (Dana et al., 2016) in PG cells using a Cre-dependent viral vector (Fig. 7A). Separate excitation lasers and selective emission filters (see Methods) were used to simultaneously and selectively image from Thy1+ sTCs and GAD2+ PG cells in the same field of view or from the neuropil of the same glomerulus (Fig. 7B). Signals imaged from the glomerular neuropil were generally similar for the Thy1+ signal, which reflected summed activity in mitral as well as tufted cell primary dendrites, and the GAD2+ signal, which reflected summed activity across PG cell processes (Fig. 7C). The slower decay in the GAD2+ signal is consistent with the slower decay of jRGECO1a after a calcium transient as compared to GCaMP6f (Dana et al., 2016). However, imaging from individual PG and sTC somata associated with the same glomerulus revealed diversity in inhalation-triggered dynamics, with different cells showing distinct time to peak and response durations (Fig 7D). While the numbers of sister PG and sTC cells imaged from the same glomerulus were not sufficient for strong statistical analysis, these observations directly demonstrate that distinct inhalation-triggered temporal dynamics can emerge among different neurons innervating the same glomerulus.

**Figure 7.**
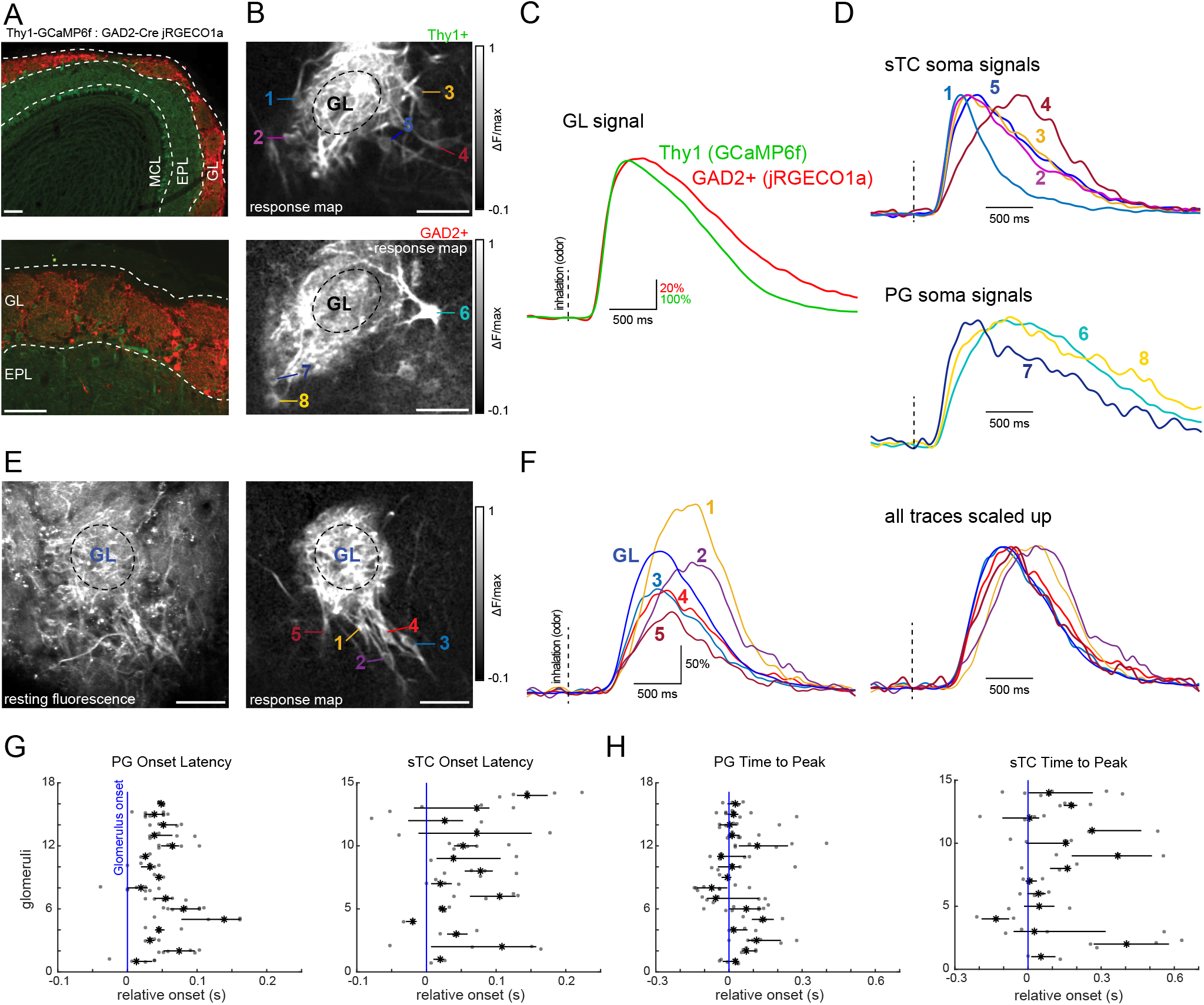
Diversity of inhalation-linked temporal dynamics is greater in sTC than PG cells innervating the same glomerulus. A. Tissue sections showing GCaMP6f expression (green) in Thy1+ MTCs (green) and jRGECO1a expression in GAD2+ juxtaglomerular neurons, after Flex.jRGECO1a (red) virus injection in a Thy1-GCaMP6f:GAD2-Cre mouse (see Text). Scale bar, 100 µm. B. Odorant-evoked ITA response map showing activation of MTCs recorded in the green channel (top) and GAD2+ neurons recorded simultaneously in the red channel (bottom). Numbers indicate presumed sTCs or PG cells whose responses are shown in (D). Odorant was ethyl butyrate. Scale bar, 50 µm. C. Overlay of ITA traces for the Thy1+ and GAD2+ signals recorded from the neuropil of the glomerulus shown in (B). D. ITA traces from the somata indicated in (B), with presumptive sTCs (top) and PG cells (bottom) overlaid with each other. All cells appear to innervate the same glomerulus. A 5 Hz low pass filter was applied to all traces prior to averaging. E. Resting fluorescence (left) and odorant-evoked ITA response map (right) taken from a Tbet-Cre mouse expressing GCaMP6f, showing a group of sister sTCs innervating the same glomerulus. Odorant was methyl valerate. Scale bar, 50 µm. F. ITA response traces from the sTCs shown in (E), along with the glomerular signal (GL). Traces are shown at right scaled to the same peak to illustrate diversity in inhalation-linked rise-times and durations. G. Dot plots showing the distribution of onset latencies for sister PG cells (left) or sTCs (right) associated with the same glomerulus, referenced to the latency measured from the neuropil of the parent glomerulus (vertical line). Each row is a different glomerulus. Black horizontal line spans the interquartile range and the black asterisk is the median. I. Same analysis in (G) but for time to peak for the same cells and glomeruli.

To more systematically compare this diversity of PG cells and sTCs, we returned to single-wavelength imaging using GCaMP6f, focusing on collecting data from sister PG (Fig. 6BC) or sTC somata (Fig. 7EF) in separate preparations. We assessed heterogeneity among sister cells of each cell type by computing the variance in ITA onset latencies and times to peak, measured relative to that of the neuropil of the ‘parent’ glomerulus (see Methods), which includes the dendrites of all sister cells (Fig. 7G, H). Onset latencies and times to peak were more variable across sister sTCs than for PG cells: mean variance in sTC onset latencies was 45 ± 9 ms (mean ± s.e.m, n= 14 glomeruli from 6 mice) compared with 19 ± 0.01 ms (n=16 glomeruli from 4 mice) for PG cells (p=0.008, unpaired t-test, Fig. 7G), consistent with a recent report (Homma et al., 2019); mean variance in time to peak (same glomeruli and cells as above) was 123 ± 20 ms for sTCs, compared with 73 ± 36 ms for PG cells (p=0.025, unpaired t-test, Fig. 7H). This result suggests that the greater variability in inhalation-triggered response dynamics observed across the population of sTCs as compared to PG cells is not an artifact of sampling across different glomeruli, but instead that this diversity can emerge within the glomerular circuit.

## DISCUSSION

A single inhalation of odorant is sufficient for odor identification, and incoming olfactory information arrives at the olfactory bulb in the form of transient bursts of OSN activity linked to each inhalation. The neural circuits that process olfactory inputs are well known, but how these circuits respond to the dynamic inputs driven by odorant inhalation in vivo remains unclear. Here we sought to better understand this key processing step by imaging from major cell types in the olfactory network and sampling odorants in the anesthetized mouse using a standard, reproducible inhalation. This approach allowed us to compare the dynamics of inhalation-linked activity as it progressed through the OB glomerular network, beginning with OSN inputs and glomerular layer interneurons thought to perform key sensory processing early in the respiratory cycle, and ending with mitral and tufted cells, which carry information out of the olfactory bulb.

Several general principles emerged. First, inhalation elicits relatively simple bursts of OSN input to a glomerulus, which occur over a limited range of latencies that is glomerulus-and odor-specific. Second, juxtaglomerular inhibitory interneurons – e.g., presumptive PG and SA cells – also show uniformly short onset latencies and simple excitatory response transients following inhalation. Third, diversity in inhalation-linked response patterns emerges at the level of glomerular output neurons, manifesting in a larger range of times to peak response and burst durations and in a higher prevalence of suppressive components of the inhalation-linked response. Finally, we find that mitral and tufted cell response patterns are highly overlapping, such that these projection neuron subtypes cannot be cleanly distinguished solely on the basis of their inhalation-linked responses. Overall, these results are consistent with a model in which diversity in inhalation-linked patterning of OB output arises first at the level of OSN inputs to the OB and is then enhanced by feedforward inhibitory circuits in the glomerular layer (Dhawale et al., 2010, Kikuta et al., 2013).

The glomerulus-and odorant-specific variation in inhalation-linked response latencies of OSN inputs is consistent with that described earlier by us and others (Spors et al., 2006, Carey et al., 2009), with latencies varying across a range of 197-321 ms (25^th^ – 75^th^ percentiles). Notably, PG and SA cells showed a near-identical distribution of response patterns, with responses overwhelmingly consisting of simple and brief inhalation-driven bursts of excitation. However, a fraction of SA cells displayed response durations that were prolonged relative to those of PG cells. In contrast, diversity in inhalation-locked mitral and tufted cell activity could not be fully accounted for by diversity in OSN inputs. Both mitral and tufted cell populations displayed a larger range of inhalation-linked onset latencies and burst durations than seen among OSNs, and inhalation-linked response patterns could include multiphasic excitatory components – features which were rare or absent among OSNs.

Our data also allowed us to compare inhalation-linked response patterns of mitral versus tufted cells. Surprisingly, at the population level, we found little difference in inhalation-linked dynamics between these two populations. However, the prevalence of clean air evoked ITA responses was greater among sTCs. Furthermore, sTCs defined by their expression of the neuropeptide transmitter CCK did show a significantly shorter range of response latencies and durations than mitral cells or the wider population of sTCs. However, the distribution of sTC and mitral cell response patterns overlapped a great deal: mitral and sTCs (as defined by soma location regardless of genetic marker) were not significantly different in mean onset latencies, and the mode of their latency distribution was identical for the two cell types. Finally, while relatively rare compared to earlier reports (Kollo et al., 2014, Economo et al., 2016, Diaz-Quesada et al., 2018), inhalation-linked suppression was seen in both mitral cells and sTCs. Overall, our data are consistent with the long-held notion that mitral and tufted cells constitute functionally distinct subpopulations of output neurons, but indicate that these cell types cannot be distinguished solely on the basis of their inhalation-linked responses. Instead, our data indicate that the representation of olfactory information by these subpopulations, at least with respect to respiratory patterning, is highly overlapping.

Respiratory patterning of ongoing activity in the absence of odorant stimulation is well-documented and has been hypothesized, among other functions, to serve as a reference for a timing-based code for odor identity (Kepecs et al., 2006, Spors et al., 2006, Cury and Uchida, 2010, Shusterman et al., 2011, Wachowiak, 2011). Here, we observed inhalation-linked patterning of activity during inhalation of clean air in all cell types, although the prevalence of such responses (<10% across all populations, and < 5% across OSN glomeruli) was smaller than reported in earlier studies (Fukunaga et al., 2014, Diaz-Quesada et al., 2018). Indeed, in this study, excitation in response to inhalation was unique to sTCs and not observed among MCs. This lower prevalence may be a result of our use of cleaned air rather than ambient room air as our background condition, or may reflect limitations in the sensitivity and temporal resolution of the GCaMP imaging approach.

Surprisingly, a small fraction of clean air-driven responses were suppressive, with such suppression observed in sTCs, MCs and even some OSN inputs. To our knowledge, this is the first report of inhalation alone driving suppression of activity in these cell types. One explanation for these results could lie in the low frequency (0.25 Hz) of artificial inhalation used in our experiments: if some OSN populations are sensitive to odor components arising from within the animal’s own nasal cavity – for example from metabolic processes-these components could drive basal activity of OSNs and in sTCs of their target glomeruli, which would be transiently removed by each inhalation of clean air. This effect could be even more pronounced in the intact, awake mouse where OSNs are exposed to exhaled air containing metabolic odorants (Munger et al., 2010, Mori et al., 2014).

What can we infer from these comparisons about the primary synaptic interactions shaping inhalation-linked patterning of olfactory bulb output? First, the data suggest that PG and SA cell excitation largely follows OSN input dynamics, consistent with evidence from slice studies that these cells are highly sensitive to OSN stimulation mediated either by mono-or disynaptic excitation (Gire and Schoppa, 2009, Shao et al., 2009, Kiyokage et al., 2010, Najac et al., 2015). We saw little to no evidence of delayed PG/SA cell responses that would correspond to feedback excitation from the late-phase responses observed in some MTCs. In fact, only 2.3% of PG and 7.4% of SA interneuron onset latencies followed the mean M/sTC onset latency. This result is somewhat surprising, as mitral cells can mediate feedback excitation of PG and SA cells via dendrodendritic synapses in the glomerular neuropil (Najac et al., 2015). Second, our results suggest that rapid feedforward inhibition from PG or SA cells may underlie the longer-latency responses seen in some mitral and tufted cells – for example, the slowest quartile of MT responses show onset latencies that roughly match the median peak time of the PG cell response. Third, the presence of longer-duration mitral cell excitatory responses that outlast those of any OSN input suggests an additional source of excitatory drive onto mitral cells, the identity of which remains unclear. A disinhibitory circuit is unlikely, as we did not observe suppression of PG or SA cells, therefore these results suggest that the intrinsic properties of MT cells could give rise to prolonged spike bursts, or that secondary excitation by ET cells could extend the duration of M/sTC excitatory responses (Carlson et al., 2000). This prolonged excitatory component was often longer than the 4-second interval between sniffs, implying that it may be important in shaping tonic levels of excitability across multiple inhalations in the awake animal (Diaz-Quesada et al., 2018).

Overall, these results establish a basic framework for how glomerular circuits are engaged to shape inhalation-linked patterning of olfactory bulb output, with feedforward inhibitory circuits adding to the initial diversity of temporal patterns of input relayed by olfactory sensory neurons. Further experiments are necessary to integrate other olfactory bulb cell types into this framework – for example, granule cells, deep short axon cells, external plexiform layer interneurons and centrifugal inputs from the olfactory cortex may also contribute to shaping respiratory patterning of olfactory bulb output. Understanding the response dynamics of each of these cell types with respect to a single inhalation of odorant should allow for a dynamic model of olfactory bulb network function across the fundamental unit of information sampling in the olfactory system. Such a model may then be used to yield insights into olfactory processing across the full range of sampling frequencies used in the behaving animal.

## ACKNOWLEDGMENTS

This work was supported by the National Institute on Deafness and Other Communication Disorders [F32DC016531 to S.M.S, R01DC006441 to M.W.]. We thank Jackson Ball, Gustavo A. Vasquez-Opazo and Rebecca L. Kummer for excellent technical assistance, Thomas Rust for data analysis software and Tom P. Eiting, Shawn D. Burton, Isaac A. Youngstrom, Yusuke Tsuno, and Andrew K. Moran for helpful feedback and discussion.

